## Supplementary file for "DENND6A couples Arl8b to a Rab34/RILP/dynein complex regulating retrograde lysosomal trafficking and autophagy"

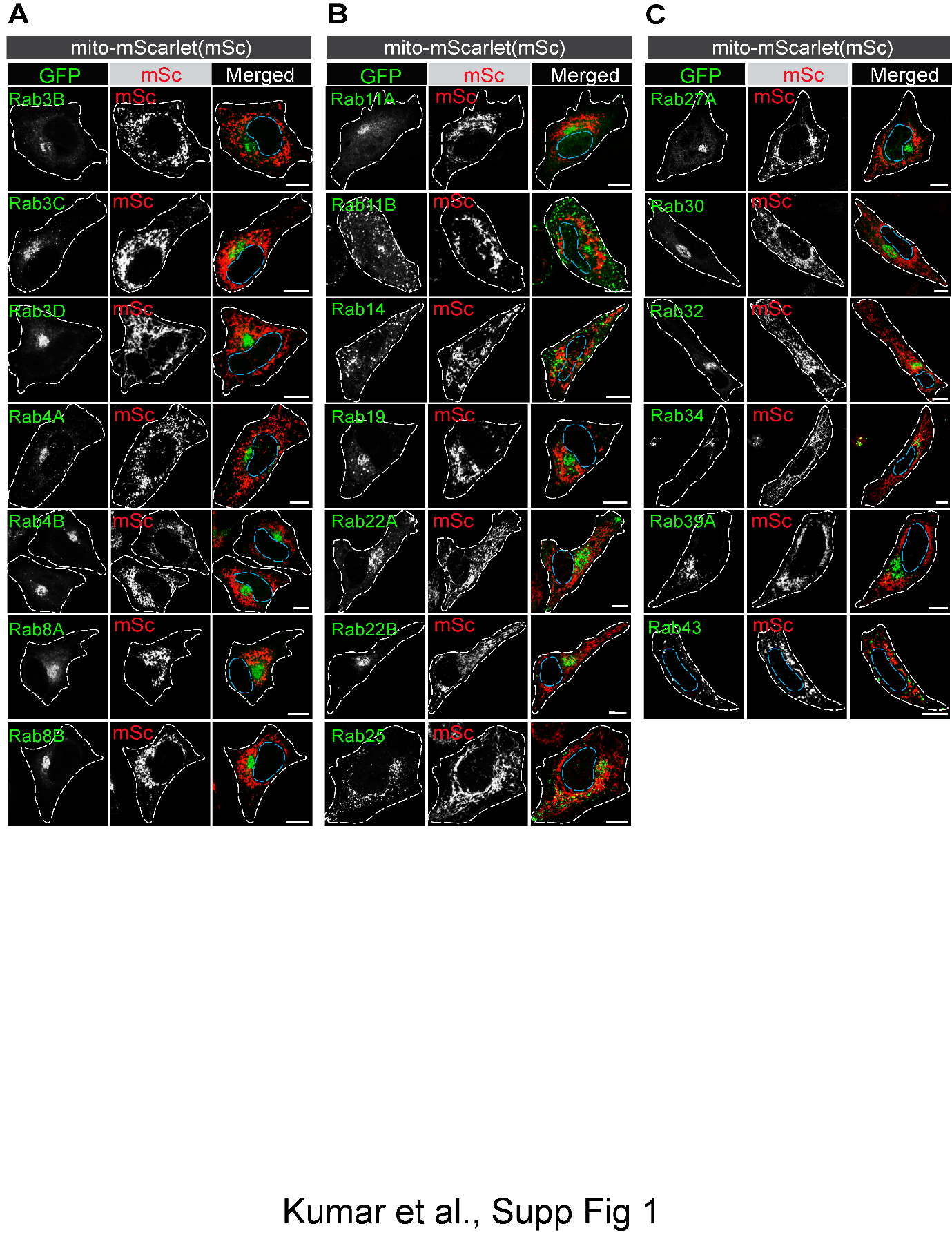

**Supplementary figure 1: mScarlet(mSc) targeted to the mitochondria does not recruit Rab GTPases. (A-C)** HeLa cells co-transfected with GFP-Rabs and mito-mSc were fixed and imaged. The nucleus and cell periphery are outlined by blue and white dotted line respectively. Scale bar = 10 µm.

**Movie 1: 3D reconstruction of SIM imaging of fixed HeLa cells expressing DENND6A-GFP and stained with LAMP1 antibody.** HeLa cells expressing DENND6A-GFP were fixed, stained with LAMP1 antibody. 3D-SIM images were acquired using LSM880-Elyra PS1 super-resolution microscopy. The scale bar is indicated at the bottom left corner. DENND6A-GFP is indicated in blue and LAMP1 is indicated in red.

**Movie 2: 3D reconstruction of SIM imaging of fixed HeLa cells expressing GFP and stained with LAMP1 antibody.** HeLa cells expressing GFP alone fixed, stained with LAMP1 antibody. 3D-SIM images were acquired using LSM880-Elyra PS1 super-resolution microscopy. The scale bar is indicated at the bottom left corner. GFP is indicated in blue and LAMP1 is indicated in red.
